# Association of *Prevotella* enterotype with polysomnographic data in obstructive sleep apnea/hypopnea syndrome patients

**DOI:** 10.1101/394064

**Authors:** Chih-Yuan Ko, Ji-Mim Fan, An-Ke Hu, Li-Mei Huang, Huan-Zhang Su, Jiao-Hong Yang, Hua-Ping Zhang, Yi-Ming Zeng

## Abstract

Intermittent hypoxia and sleep fragmentation are critical pathophysiological processes involved in obstructive sleep apnea/hypopnea syndrome (OSAHS). These manifestation independently affect similar brain regions and contribute to OSAHS-related comorbidities that are known to be related to the host gut alteration microbiota. We hypothesized that microbiota disruption influences the pathophysiological processes of OSAHS through a microbiota–gut–brain axis. Thus, we aim to survey enterotypes and polysomnographic data of OSAHS patients. Subjects were diagnosed by polysomnography, from whom fecal samples were obtained and analyzed for the microbiome composition by variable regions 3–4 of 16S rRNA pyrosequencing and bioinformatic analyses. We examined blood cytokines level of all subjects. Three enterotypes *Bacteroides* (n=73), *Ruminococcus* (n=14), and *Prevotella* (n=26) were identified. Central apnea indices, mixed apnea indices, N1 sleep stage, mean apnea–hypopnea duration, and arousal indices were increased in apnea–hypopnea indices (AHI) ≥15 patients with the *Prevotella* enterotype. However, for AHI<15 subjects, obstructive apnea indices and systolic blood pressure were significantly observed in *Ruminococcus* and *Prevotella* enterotypes, respectively. The present study indicates the possibility of pathophysiological interplay between enterotypes and sleep structure disruption in sleep apnea through a microbiota–gut–brain axis and offers some new insight toward the pathogenesis of OSAHS.

**Importance:** Intermittent hypoxia (IH) and sleep fragmentation (SF) are hallmarks of are the predominant mechanism underlying obstructive sleep apnea/hypopnea syndrome (OSAHS). Moreover, IH and SF of pathophysiological roles in the gut microbiota dysbiosis in OSAHS have been demonstrated. We hypothesized that gut microbiota disruption may cross-talk the brain function via microbiota–gut–brain axis. Indeed, we observed central apnea indices and other parameters of disturbances during sleep were significantly elevated in AHI≥15 patients with the *Prevotella* enterotype. This enterotype prone to endotoxin production, driving systemic inflammation, ultimately contributes to OSAHS-linked comorbidities. Vice versa, increasing the arousal index leads to systemic inflammatory changes and accompanies metabolic dysfunction. We highlight that the possibility that the microbiota–gut–brain axis operates a bidirectional effect on the development of OSAHS pathology.

## Introduction

Intermittent hypoxia (IH) and sleep fragmentation (SF) are hallmarks of obstructive sleep apnea/hypopnea syndrome (OSAHS) [1-3]. IH plays a critical pathophysiological role in of OSAHS, often accompanied by reduced oxygen saturation, increased systemic pressure and bloodstream, excessive sympathetic neural activity, impairment of autonomic function and apnea episodes end with an arousal of the central nervous system (CNS), ultimately result in vascular endothelial dysfunction and multi-organ morbid consequences. The underlying mechanism involves inflammation and oxidative stress cascades [4,5].

Contrastingly, sleep structure disruption is another risk factor for the pathophysiology of OSAHS, causing major end-organ morbidity independent of IH [6,7]. Repeated arousals disturbing different stages of sleep are the predominant mechanism underlying OSAHS-induced brain injury wherein results from disruptions of rapid eye movement (REM) and non-REM (NREM) [7]. Even disturbances in sleep continuity are associated with emotional disorders [8]. SF promotes obesity and metabolic abnormalities and may be mediated by concurrent alterations of the host gut microbiota and concurrent systemic and adipose tissue inflammatory alterations accompanied by insulin resistance [3,9]. Prolongation of the N1 stage and shortening of REM times were observed in OSAHS-induced hypertension patients. Reportedly, prolongation of the N1 sleep stage causes elevation of fasting blood glucose [10]. Elevated serum lipopolysaccharide (LPS)-binding protein levels might prolong the N1 stage and increase SF, which may be related to increased nighttime respiratory events and arousals [10]. Interestingly, the disturbance of sleep structure also contributes to mild cognitive decline in OSAHS [7]. However, treating OSAHS patients with continuous positive airway pressure (CPAP) has protective effects on neurocognition, and it has been postulated that the microbiota can be modulated during CPAP treatment [11], implying that the microbiota might participate in the pathophysiological developed mechanism.

Emerging evidence suggests that the gut microbiota play a crucial role in modulating the risk of several chronic diseases and maintaining intestinal immunity and whole body homeostasis. These effects have important implications for diseases such as obesity, cardiometabolic abnormalities, inflammatory bowel disease (IBD), and mental illness [12]. Additionally, the gut microbiota alterations manifested in IH and FS mimic in OSAHS animal models [1,3]. However, some of the underlying mechanisms of OSAHS-related comorbidities remain unclear. Enterotype analysis has been proposed as a useful method to understand human gut microbial communities, including *Bacteroides, Ruminococcus*, and *Prevotella* enterotypes, irrespective of ethnicity, gender, age or body mass index (BMI) [13]. Moreover, enterotypes subdivision provides an attractive framework for linking human disease. For example, *Bacteroides* enterotype has been reported to pose an increased risk for IBD [14-16].

Notably, the characteristics of IH and SF in OSAHS can trigger the inflammatory response, which then alters the intestinal microbial community composition [1,3,9]. Conversely, gut microbiomes can also respond to the brain via the microbiota–gut– brain axis, as has been reported in psychiatric disorders [17,18]. However, this hypothesis has not been verified for OSAHS. Thus, the present study tested the hypothesis that the microbiota–gut–brain axis is involved in the pathogenesis of OSAHS. We examined whether impaired sleep architecture is associated with gut microbiota alteration by investigating sleep parameters of polysomnography (PSG) data and pro-inflammatory cytokines in various enterotypes of OSAHS subjects.

## Results

### Patient characteristics and enterotype distribution

We enrolled 113 patients (61, AHI<15; 52, AHI≥15). Patients were divided according to three enterotypes: *Bacteroides* (n=73), *Ruminococcus* (n=14), and *Prevotella* (n=26) (Figure 1). The ages of the *Ruminococcus* enterotype patients were significantly higher than those of the *Bacteroides* enterotype patients (Table 1). BMI and hip circumference of the *Prevotella* enterotype patients were significantly higher than those of the *Bacteroides* enterotype patients (Table 1).

**Figure 1.**
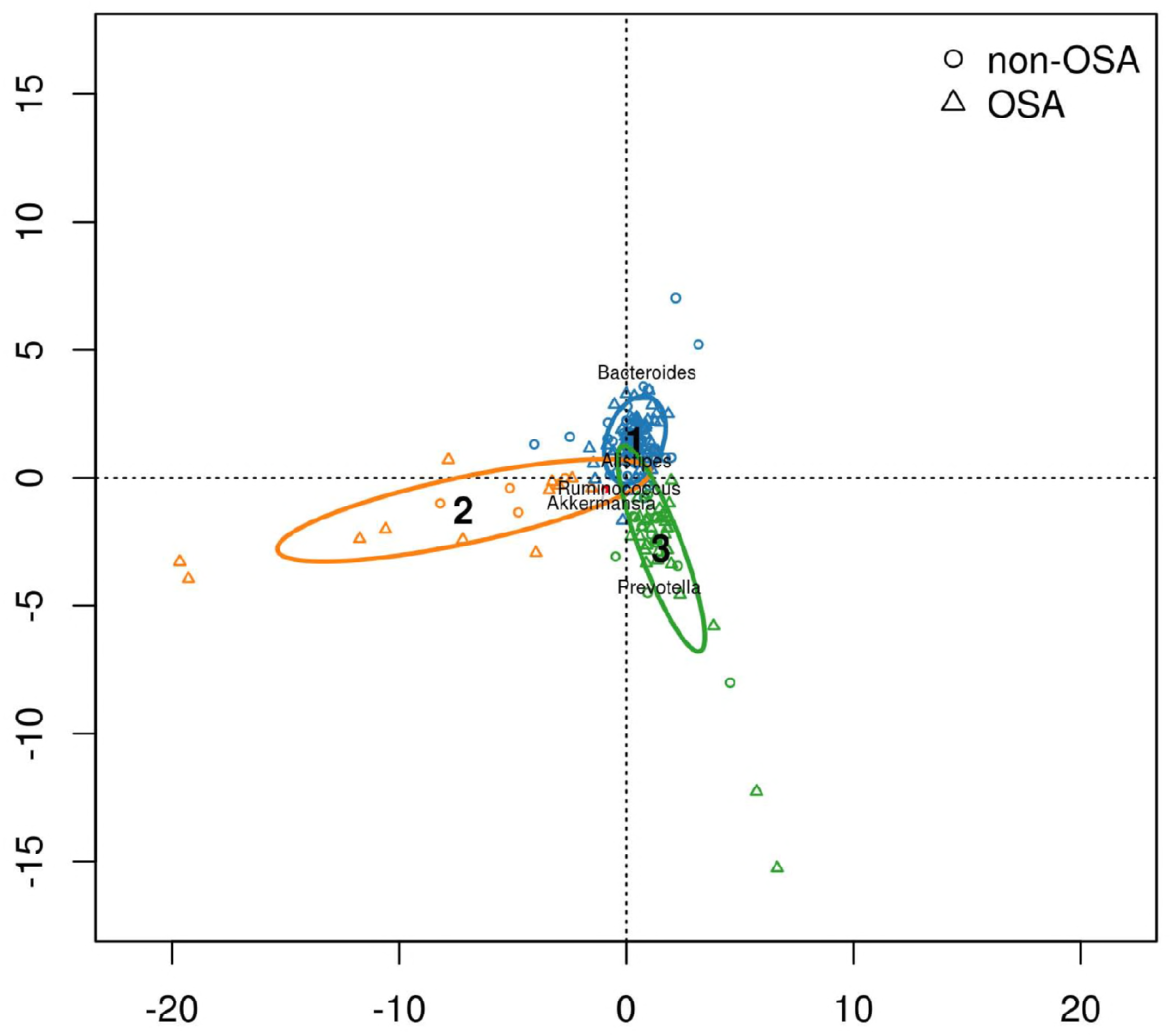
The faecal taxa of non-obstructive sleep apnoea–hypopnea syndrome (OSAHS) and OSAHS subjects of three enterotypes. Apnoea–hypopnea indices (AHI) < 15 as non-OSAHS, AHI ≥ 15 as OSAHS. Enterotype 1: *Bacteroides*, Enterotype 2: *Ruminococcus*, Enterotype 3: *Prevotella*.

**Table 1.** Participant characteristics.

|  | *Bacteroides*<br>enterotype<br>(n=73) | *Ruminococcus*<br>enterotype<br>(n=14) | *Prevotella*<br>enterotype<br>(n=26) |
| --- | --- | --- | --- |
| Gender (male/female) | 61/12 | 11/3 | 20/6 |
| Age (years, mean ± SD) | **40.89±10.56b** | **53.14±14.37a** | **44.58±13.51ab** |
| Height (cm) | 166.28±6.90a | 164.61±7.30a | 166.83±8.83a |
| Weight (kg) | 72.51±14.31a | 71.62±14.25a | 79.41±13.70a |
| Body mass index (kg m<sup>-2</sup>) | **26.11±3.87b** | **26.44±5.50ab** | **28.73±5.27a** |
| Waist circumference (cm) | 90.77±9.87a | 91.21±13.33a | 96.54±12.27a |
| Hip circumference (cm) | **96.95±6.03b** | **97.75±10.42ab** | **103.21±8.97a** |
| Waist-to-hip ratio | 0.94±0.07a | 0.93±0.06a | 0.93±0.06a |

### PSG parameter analysis

Comparisons among patients with different enterotypes showed that central apnea times, mixed apnea index, and mixed apnea times were the highest in the *Prevotella* enterotype patients (Table 2).

**Table 2.** Polysomnographic data analysis in three enterotypes subjects.

|  | *Bacteroides*<br>enterotype<br>(n=73) | *Ruminococcus*<br>enterotype<br>(n=14) | *Prevotella*<br>enterotype<br>(n=26) |
| --- | --- | --- | --- |
| Total sleep time (min) | 389.01±99.14a | 362.77±120.12a | 429.64±119.1a |
| N1 sleep stage (min) | 138.56±66.92a | 152.04±82.37a | 171.62±96.22a |
| N1 sleep stage (%) | 36.10±15.90a | 42.19±14.54a | 39.76±17.28a |
| N2 sleep stage (min) | 111.30±60.67a | 85.85±44.25a | 111.94±57.67a |
| N2 sleep stage (%) | 28.10±11.99a | 24.45±9.62a | 25.86±10.48a |
| N3 sleep stage (min) | 71.91±36.85a | 57.71±45.99a | 66.44±41.66a |
| N3 sleep stage (%) | 18.33±8.29a | 15.95±10.23a | 16.25±9.79a |
| Non-rapid-eye movement (NREM) (min) | 321.79±87.88a | 295.60±96.74a | 350.01±98.5a |
| NREM (%) | 82.56±10.84a | 82.57±7.62a | 81.87±9.88a |
| Rapid-eye movement (REM) (min) | 67.21±43.27a | 67.17±37.26a | 79.63±44.02a |
| REM (%) | 17.43±10.84a | 17.42±7.62a | 18.12±9.88a |
| Sleep efficiency (%) | 70.40±14.99a | 64.62±21.45a | 75.92±18.36a |
| Sleep latency | 31.0±29.96a | 31.70±29.97a | 18.93±20.36a |
| Wake after sleep onset | 129.91±71.69a | 174.76±116.12a | 110.50±98.33a |
| Arousal time (min) | 157.64±78.71a | 202.23±131.03a | 128.14±100.46a |
| Arousal times | 20.35±11.57a | 19.64±12.70a | 14.50±9.36a |
| Arousal index (events/h) | 3.34±2.06a | 3.86±3.19a | 2.30±1.68a |
| Arousal time in REM | 39.93±38.94a | 39.42±34.74a | 52.00±36.13a |
| Arousal index in REM | 32.25±16.44a | 29.52±17.66a | 38.96±16.29a |
| Arousal time in NREM | 210.15±126.03a | 209.57±148.79a | 288.19±226.67a |
| Arousal index in NREM | 38.17±15.54a | 39.98±16.20a | 45.76±26.09a |
| Total sleep arousal times | 250.08±136.58a | 249.00±166.24a | 340.19±249.89a |
| Total sleep arousal index | 38.06±15.18a | 38.75±15.83a | 44.64±24.15a |
| Apnea-hypopnea index (events/h) | 19.63±21.42a | 27.90±22.63a | 27.41±30.96a |
| Apnea-hypopnea times | 132.19±170.66a | 180.50±176.90a | 221.34±278.37a |
| Obstructive apnea index (events/h) | 8.85±15.04a | 10.88±12.44a | 10.62±16.09a |
| Obstructive apnea times | 62.78±120.68a | 71.28±102.99a | 85.26±134.75a |
| Central apnea index (events/h) | 0.46±1.29a | 2.75±8.64a | 1.92±4.04a |
| Central apnea times | **3.39±9.45b** | **11.78±31.62ab** | **17.5±40.21a** |
| Mixed apnea index (events/h) | **0.59±1.52b** | **1.61±3.80ab** | **2.88±6.79a** |
| Mixed apnea times | **4.56±13.25b** | **10.5±27.95ab** | **25.73±64.8a** |
| Hypopnea index (events/h) | 9.71±8.95a | 12.63±17.71a | 11.98±14.29a |
| Hypopnea times | 61.45±55.31a | 86.92±136.55a | 92.84±125.09a |
| Longest apnea time (s) | 37.80±23.24a | 46.64±27.65a | 45.26±38.39a |
| Mean apnea-hypopnea duration (s) | 19.55±8.98a | 20.25±8.28a | 20.01±9.78a |
| Longest hypopnea time (s) | 68.26±33.06a | 66.85±27.29a | 78.50±32.89a |
| Average hypopnea time (s) | 28.30±9.95a | 29.75±12.62a | 31.52±8.38a |
| Oxygen desaturation index (events/h) | 18.72±20.79a | 25.22±23.56a | 25.9±29.82a |
| Lowest oxygen saturation (%) | 82.26±8.81a | 79.35±9.34a | 82.23±10.88a |
| Average oxygen saturation (%) | 94.34±2.04a | 93.92±3.34a | 94.15±2.66a |
| Longest oxygen desaturation (s) | 109.16±42.98a | 95.17±28.61a | 99.51±40.54a |
| Mean heart rate | 65.95±10.32a | 65.00±7.93a | 63.73±9.55a |
| Arrhythmia index (events/h) | 12.89±54.36a | 38.04±134.47a | 4.23±8.19a |
| Maximum heart rate | 99.41±15.06a | 97.28±12.19a | 96.92±16.70a |
| Minimum heart rate | 51.67±8.97a | 53.28±5.21a | 51.53±6.68a |
| Blood pressure elevation index (events/h) | 13.22±15.21a | 16.43±15.52a | 16.58±22.43a |
| The highest systolic blood pressure (mmHg) | 157.98±39.08a | 148.57±58.14a | 158.80±45.51a |
| Average systolic blood pressure (mmHg) | 120.41±25.98a | 113.78±42.26a | 118.19±28.64a |
| Average diastolic blood pressure (mmHg) | 81.80±16.05a | 77.21±24.46a | 82.92±21.59a |

Using 15 as the AHI cut-off, when AHI≥15, N1 sleep stage, arousal time in REM, arousal index in REM, arousal time in NREM, total sleep arousal times, total sleep arousal index, central apnea times, mixed apnea index, and mixed apnea times were the highest in the *Prevotella* enterotype patients. Contrastingly, sleep latency and arousal time were the lowest in the *Prevotella* enterotype patients (Table 3). When AHI<15, obstructive apnea index and obstructive apnea times were the highest in the *Ruminococcus* enterotype subjects, the highest and average systolic BP were the highest in the *Prevotella* enterotype subjects (Table 4).

**Table 3.**
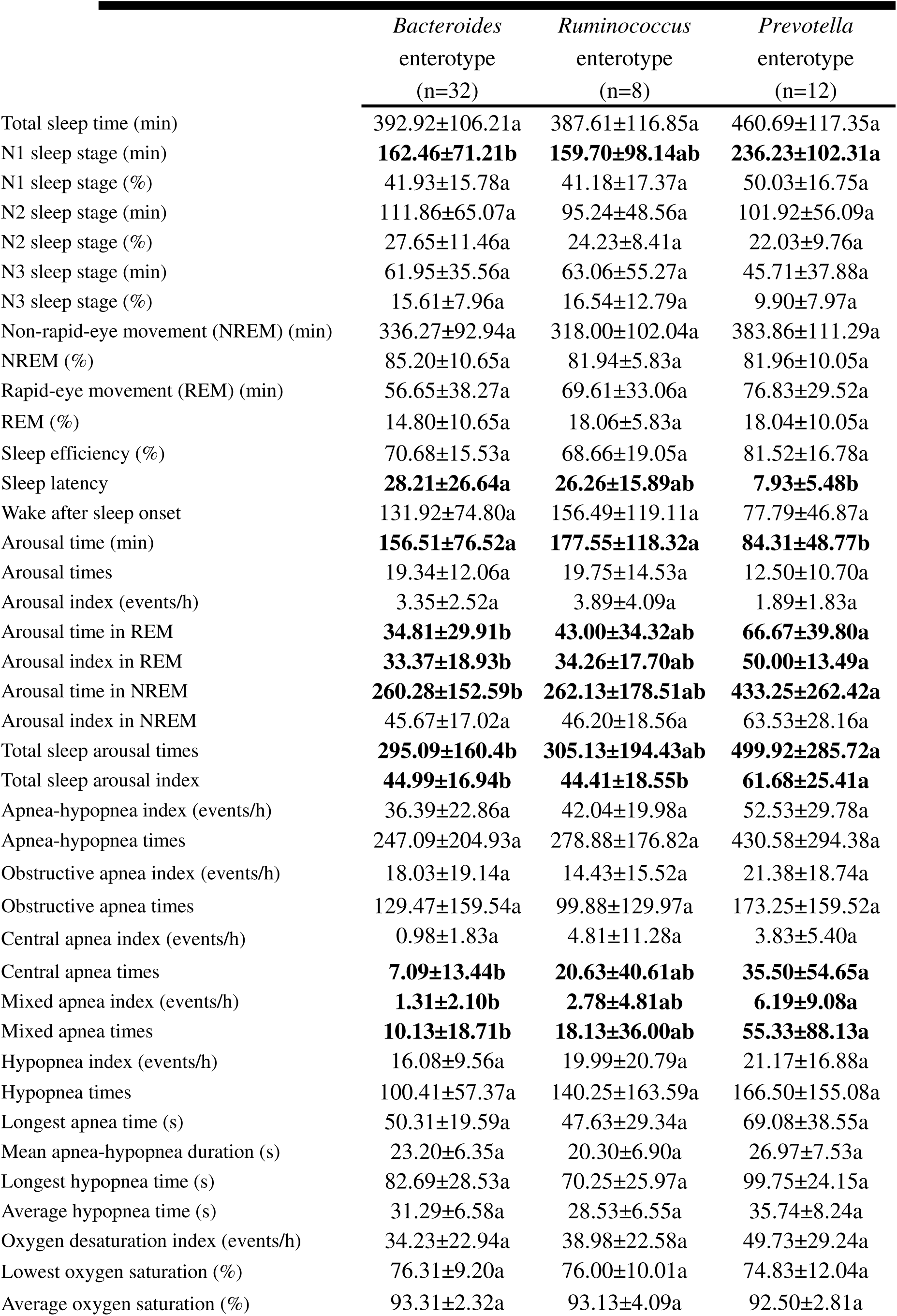
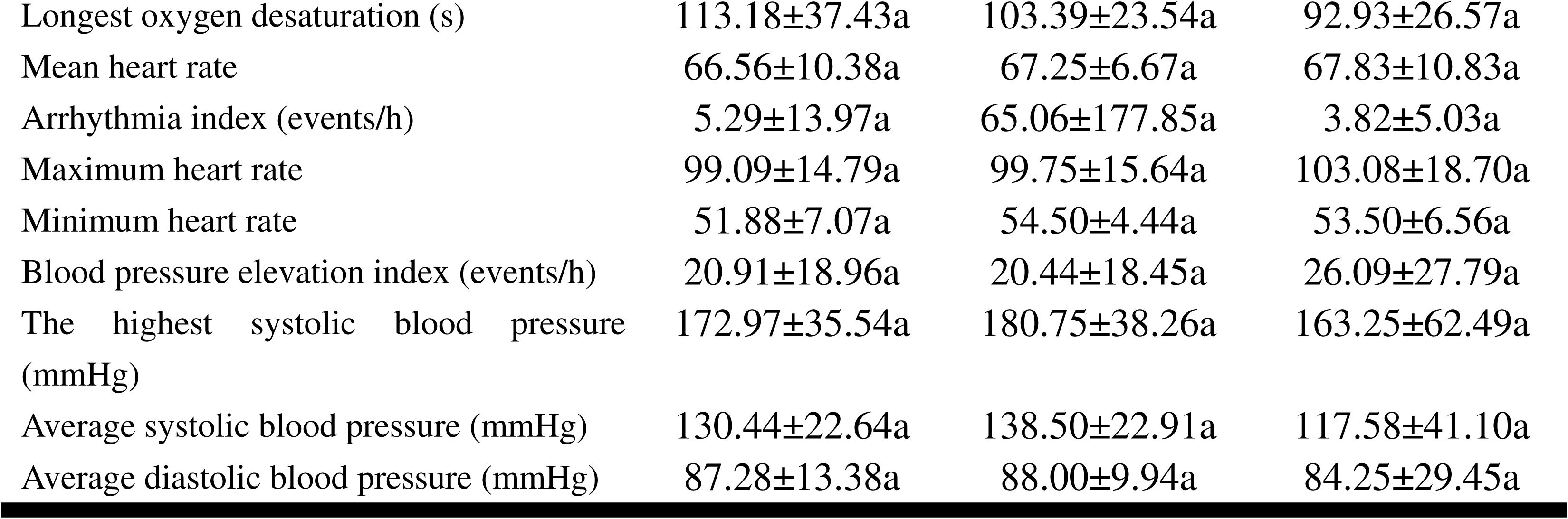
Polysomnographic data analysis in three enterotypes of apnoea– hypopnea indices ≥ 15 patients.

|  | *Bacteroides*<br>enterotype<br>(n=32) | *Ruminococcus*<br>enterotype<br>(n=8) | *Prevotella*<br>enterotype<br>(n=12) |
| --- | --- | --- | --- |
| Total sleep time (min) | 392.92±106.21a | 387.61±116.85a | 460.69±117.35a |
| N1 sleep stage (min) | **162.46±71.21b** | **159.70±98.14ab** | **236.23±102.31a** |
| N1 sleep stage (%) | 41.93±15.78a | 41.18±17.37a | 50.03±16.75a |
| N2 sleep stage (min) | 111.86±65.07a | 95.24±48.56a | 101.92±56.09a |
| N2 sleep stage (%) | 27.65±11.46a | 24.23±8.41a | 22.03±9.76a |
| N3 sleep stage (min) | 61.95±35.56a | 63.06±55.27a | 45.71±37.88a |
| N3 sleep stage (%) | 15.61±7.96a | 16.54±12.79a | 9.90±7.97a |
| Non-rapid-eye movement (NREM) (min) | 336.27±92.94a | 318.00±102.04a | 383.86±111.29a |
| NREM (%) | 85.20±10.65a | 81.94±5.83a | 81.96±10.05a |
| Rapid-eye movement (REM) (min) | 56.65±38.27a | 69.61±33.06a | 76.83±29.52a |
| REM (%) | 14.80±10.65a | 18.06±5.83a | 18.04±10.05a |
| Sleep efficiency (%) | 70.68±15.53a | 68.66±19.05a | 81.52±16.78a |
| Sleep latency | **28.21±26.64a** | **26.26±15.89ab** | **7.93±5.48b** |
| Wake after sleep onset | 131.92±74.80a | 156.49±119.11a | 77.79±46.87a |
| Arousal time (min) | **156.51±76.52a** | **177.55±118.32a** | **84.31±48.77b** |
| Arousal times | 19.34±12.06a | 19.75±14.53a | 12.50±10.70a |
| Arousal index (events/h) | 3.35±2.52a | 3.89±4.09a | 1.89±1.83a |
| Arousal time in REM | **34.81±29.91b** | **43.00±34.32ab** | **66.67±39.80a** |
| Arousal index in REM | **33.37±18.93b** | **34.26±17.70ab** | **50.00±13.49a** |
| Arousal time in NREM | **260.28±152.59b** | **262.13±178.51ab** | **433.25±262.42a** |
| Arousal index in NREM | 45.67±17.02a | 46.20±18.56a | 63.53±28.16a |
| Total sleep arousal times | **295.09±160.4b** | **305.13±194.43ab** | **499.92±285.72a** |
| Total sleep arousal index | **44.99±16.94b** | **44.41±18.55b** | **61.68±25.41a** |
| Apnea-hypopnea index (events/h) | 36.39±22.86a | 42.04±19.98a | 52.53±29.78a |
| Apnea-hypopnea times | 247.09±204.93a | 278.88±176.82a | 430.58±294.38a |
| Obstructive apnea index (events/h) | 18.03±19.14a | 14.43±15.52a | 21.38±18.74a |
| Obstructive apnea times | 129.47±159.54a | 99.88±129.97a | 173.25±159.52a |
| Central apnea index (events/h) | 0.98±1.83a | 4.81±11.28a | 3.83±5.40a |
| Central apnea times | **7.09±13.44b** | **20.63±40.61ab** | **35.50±54.65a** |
| Mixed apnea index (events/h) | **1.31±2.10b** | **2.78±4.81ab** | **6.19±9.08a** |
| Mixed apnea times | **10.13±18.71b** | **18.13±36.00ab** | **55.33±88.13a** |
| Hypopnea index (events/h) | 16.08±9.56a | 19.99±20.79a | 21.17±16.88a |
| Hypopnea times | 100.41±57.37a | 140.25±163.59a | 166.50±155.08a |
| Longest apnea time (s) | 50.31±19.59a | 47.63±29.34a | 69.08±38.55a |
| Mean apnea-hypopnea duration (s) | 23.20±6.35a | 20.30±6.90a | 26.97±7.53a |
| Longest hypopnea time (s) | 82.69±28.53a | 70.25±25.97a | 99.75±24.15a |
| Average hypopnea time (s) | 31.29±6.58a | 28.53±6.55a | 35.74±8.24a |
| Oxygen desaturation index (events/h) | 34.23±22.94a | 38.98±22.58a | 49.73±29.24a |
| Lowest oxygen saturation (%) | 76.31±9.20a | 76.00±10.01a | 74.83±12.04a |
| Average oxygen saturation (%) | 93.31±2.32a | 93.13±4.09a | 92.50±2.81a |
| Longest oxygen desaturation (s) | 113.18±37.43a | 103.39±23.54a | 92.93±26.57a |
| Mean heart rate | 66.56±10.38a | 67.25±6.67a | 67.83±10.83a |
| Arrhythmia index (events/h) | 5.29±13.97a | 65.06±177.85a | 3.82±5.03a |
| Maximum heart rate | 99.09±14.79a | 99.75±15.64a | 103.08±18.70a |
| Minimum heart rate | 51.88±7.07a | 54.50±4.44a | 53.50±6.56a |
| Blood pressure elevation index (events/h) | 20.91±18.96a | 20.44±18.45a | 26.09±27.79a |
| The highest systolic blood pressure (mmHg) | 172.97±35.54a | 180.75±38.26a | 163.25±62.49a |
| Average systolic blood pressure (mmHg) | 130.44±22.64a | 138.50±22.91a | 117.58±41.10a |
| Average diastolic blood pressure (mmHg) | 87.28±13.38a | 88.00±9.94a | 84.25±29.45a |

**Table 4.** Polysomnographic data analysis in enterotypes of apnoea–hypopnea indices < 15 subjects.

|  | *Bacteroides*<br>enterotype<br>(n=41) | *Ruminococcus*<br>enterotype<br>(n=6) | *Prevotella*<br>enterotype<br>(n=14) |
| --- | --- | --- | --- |
| Total sleep time (min) | 385.96±94.49a | 329.65±126.87a | 403.04±118.21a |
| N1 sleep stage (min) | 119.91±57.60a | 141.83±62.74a | 116.25±42.34a |
| N1 sleep stage (%) | 31.55±14.61a | 43.55±11.10a | 30.96±12.49a |
| N2 sleep stage (min) | 110.88±57.82a | 73.33±38.22a | 120.54±59.68a |
| N2 sleep stage (%) | 28.47±12.52a | 24.77±11.88a | 29.15±10.26a |
| N3 sleep stage (min) | 79.70±36.37a | 50.58±33.42a | 84.21±37.23a |
| N3 sleep stage (%) | 20.47±8.01a | 15.17±6.46a | 21.70±7.83a |
| Non-rapid-eye movement (NREM) (min) | 310.49±83.12a | 265.73±88.78a | 321.00±78.85a |
| NREM (%) | 80.51±10.68a | 83.42±10.10a | 81.80±10.11a |
| Rapid-eye movement (REM) (min) | 75.47±45.56a | 63.92±45.35a | 82.04±54.54a |
| REM (%) | 19.49±10.67a | 16.58±10.10a | 18.20±10.11a |
| Sleep efficiency (%) | 70.20±14.75a | 59.23±25.03a | 71.13±18.87a |
| Sleep latency | 33.20±32.47a | 38.95±43.26a | 28.36±23.75a |
| Wake after sleep onset | 128.36±70.05a | 199.13±118.10a | 138.54±122.07a |
| Arousal time (min) | 158.52±81.32a | 235.15±150.88a | 165.71±118.75a |
| Arousal times | 21.15±11.26a | 19.50±11.13a | 16.21±8.06a |
| Arousal index (events/h) | 3.35±1.66a | 3.83±1.75a | 2.66±1.52a |
| Arousal time in REM | 43.93±44.70a | 34.67±37.97a | 39.43±28.30a |
| Arousal index in REM | 31.39±14.39a | 23.20±16.99a | 29.50±12.13a |
| Arousal time in NREM | 171.02±83.45a | 139.50±51.31a | 163.86±66.82a |
| Arousal index in NREM | 32.33±11.41a | 31.70±7.47a | 30.54±9.82a |
| Total sleep arousal times | 214.95±103.70a | 174.17±84.67a | 203.29±85.31a |
| Total sleep arousal index | 32.66±11.13a | 31.20±7.07a | 30.04±8.81a |
| Apnea-hypopnea index (events/h) | 6.56±4.54a | 9.07±5.20a | 5.88±3.43a |
| Apnea-hypopnea times | 42.51±32.76a | 49.33±38.24a | 42.00±25.31a |
| Obstructive apnea index (events/h) | **1.70±1.93b** | **6.17±4.27a** | **1.41±1.65b** |
| Obstructive apnea times | **10.73±12.29b** | **33.17±29.64a** | **9.86±10.79b** |
| Central apnea index (events/h) | 0.07±0.15a | 0.00±0.00a | 0.30±0.77a |
| Central apnea times | 0.51±1.14a | 0.00±0.00a | 2.07±5.14a |
| Mixed apnea index (events/h) | 0.03±0.11a | 0.07±0.10a | 0.05±0.09a |
| Mixed apnea times | 0.22±0.72a | 0.33±0.52a | 0.36±0.63a |
| Hypopnea index (events/h) | 4.75±3.94a | 2.83±2.95a | 4.11±2.65a |
| Hypopnea times | 31.05±28.19a | 15.83±20.05a | 29.71±21.00a |
| Longest apnea time (s) | 28.05±21.26a | 45.33±27.91a | 24.86±24.59a |
| Mean apnea-hypopnea duration (s) | 16.70±9.74a | 20.18±10.58a | 14.06±7.29a |
| Longest hypopnea time (s) | 57.00±32.26a | 62.33±30.81a | 60.29±28.51a |
| Average hypopnea time (s) | 25.97±11.49a | 31.40±18.67a | 27.91±6.86a |
| Oxygen desaturation index (events/h) | 6.63±5.30a | 6.90±4.88a | 5.48±3.66a |
| Lowest oxygen saturation (%) | 86.9±4.88a | 83.83±6.68a | 88.57±3.34a |
| Average oxygen saturation (%) | 95.15±1.35a | 95.00±1.79a | 95.57±1.51a |
| Longest oxygen desaturation (s) | 106.03±47.08a | 84.23±33.17a | 105.15±49.88a |
| Mean heart rate | 65.49±10.37a | 62.00±9.08a | 60.21±6.89a |
| Arrhythmia index (events/h) | 18.83±71.31a | 2.02±2.09a | 4.60±10.36a |
| Maximum heart rate | 99.66±15.44a | 94.00±4.60a | 91.64±13.25a |
| Minimum heart rate | 51.51±10.30a | 51.67±6.12a | 49.86±6.54a |
| Blood pressure elevation index (events/h) | 7.23±7.34a | 11.10±9.49a | 8.44±12.61a |
| The highest systolic blood pressure (mmHg) | **146.29±38.09ab** | **105.67±53.6b** | **155.00±25.4a** |
| Average systolic blood pressure (mmHg) | **112.59±25.97a** | **80.83±40.36b** | **118.71±12.14a** |
| Average diastolic blood pressure (mmHg) | 77.54±16.80a | 62.83±31.35a | 81.79±12.63a |

We used individual enterotypes for comparison due to the effects of *Ruminococcus* and *Prevotella* enterotypes on PSG. For *Ruminococcus* enterotype patients, AHI, apnea-hypopnea times, oxygen desaturation index, highest systolic blood pressure (BP), and average systolic BP were all significantly elevated in AHI≥15 patients (Table 5).

**Table 5.**
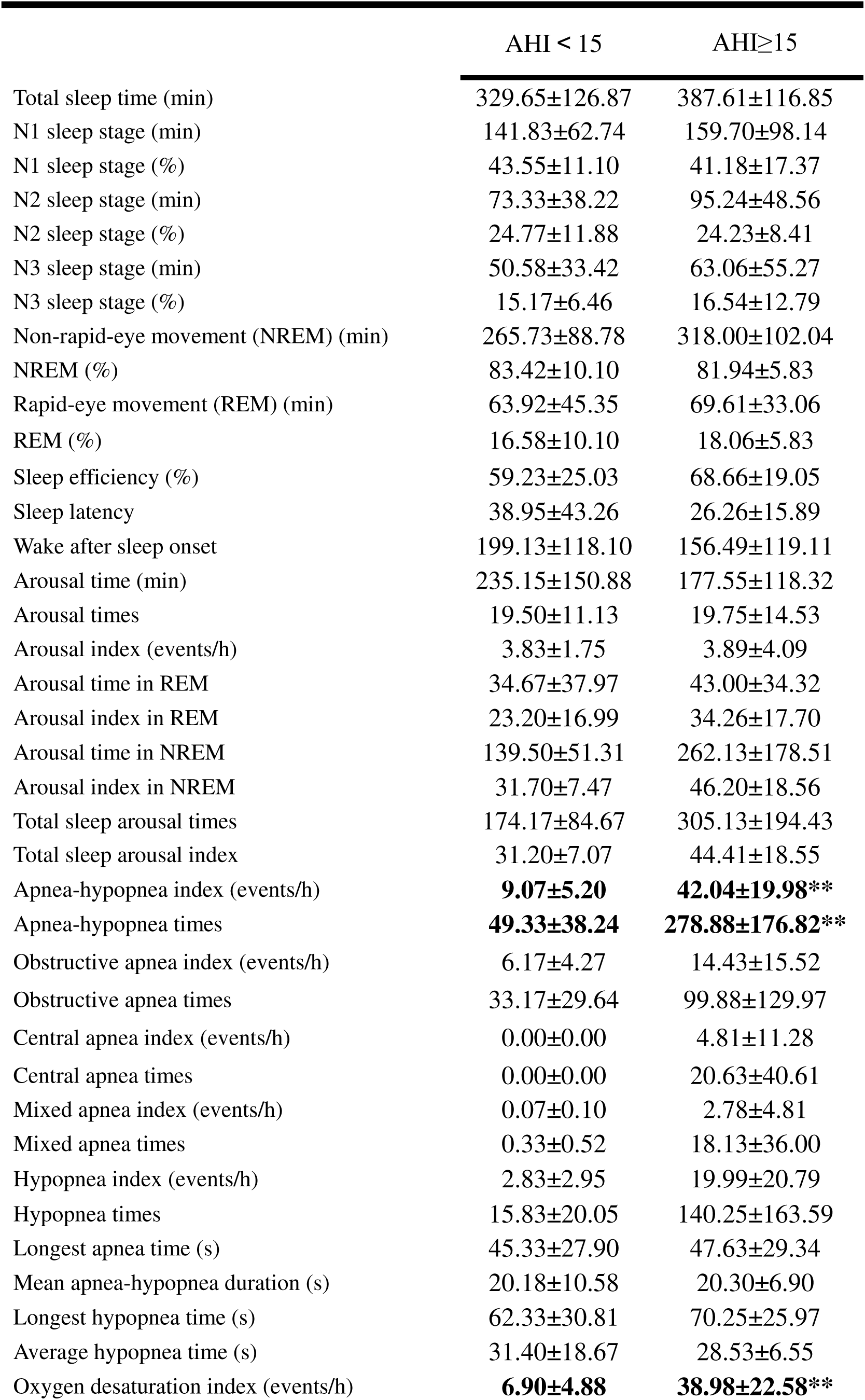
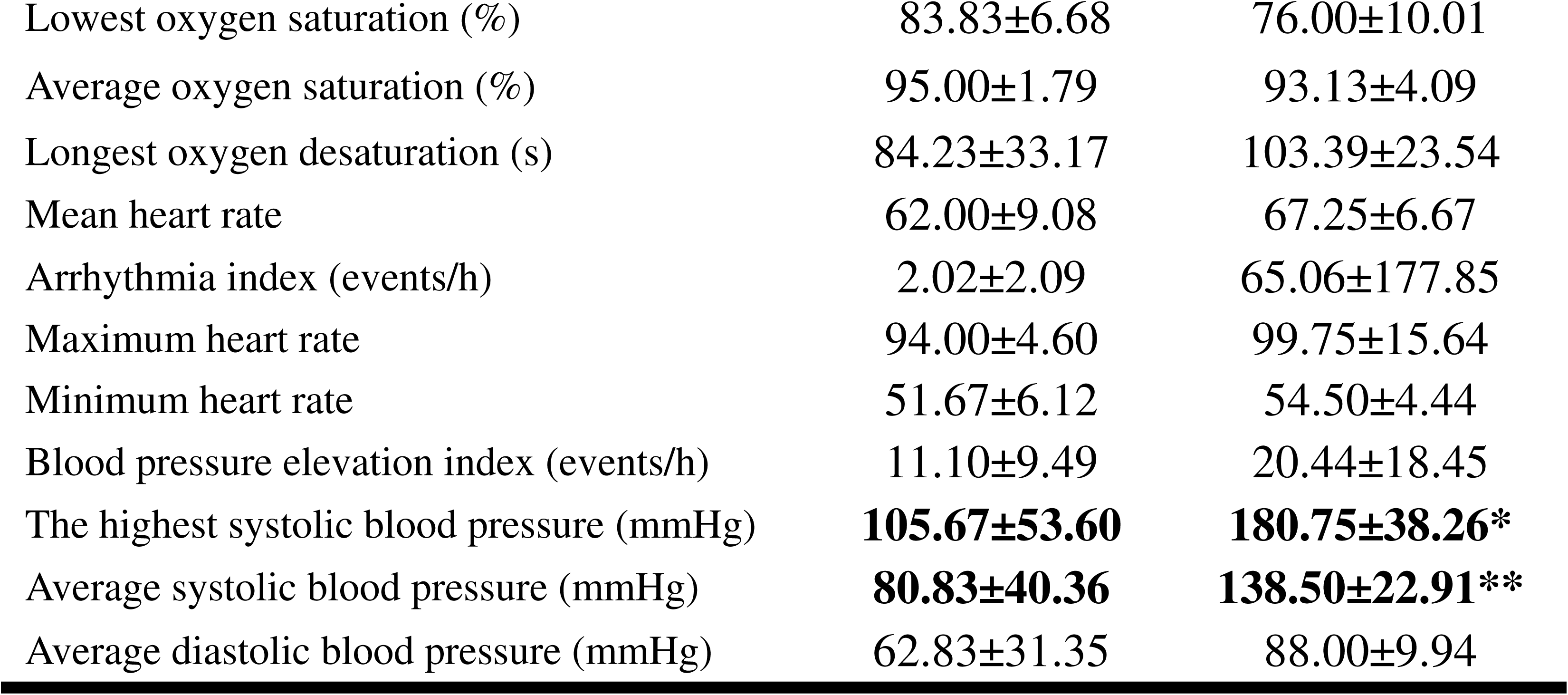
Polysomnographic data analysis in *Ruminococcus* enterotype subjects. AHI: apnoea–hypopnea indices. * p<0.05, ** p<0.01 compared with AHI<15 subjects.

For *Prevotella* enterotype patients, N1 sleep stage, N3 sleep stage, arousal index in REM, arousal time in NREM, arousal index in NREM, total sleep arousal times, total sleep arousal index, AHI, apnea-hypopnea times, obstructive apnea index, obstructive apnea times, central apnea index, mixed apnea index, hypopnea index, hypopnea times, longest apnea time, mean apnea–hypopnea duration (MAD), longest hypopnea time, average hypopnea time, oxygen desaturation index, and mean heart rate were significantly elevated in AHI≥15 patients. However, sleep latency, arousal time, lowest oxygen saturation, and average oxygen saturation were significantly decreased in AHI≥15 patients (Table 6).

**Table 6.** Polysomnographic data analysis in *Prevotella* enterotype subjects. AHI: apnoea–hypopnea indices. * p<0.05, ** p<0.01, *** p<0.001 compared with AHI<15 subjects.

|  | AHI < 15 | AHI ≥ 15 |
| --- | --- | --- |
| Total sleep time (min) | 403.04±118.21 | 460.69±117.35 |
| N1 sleep stage (min) | **116.25±42.34** | **236.23±102.31\*\*** |
| N1 sleep stage (%) | **30.96±12.49** | **50.03±16.75\*\*** |
| N2 sleep stage (min) | 120.54±59.68 | 101.92±56.09 |
| N2 sleep stage (%) | 29.15±10.26 | 22.03±9.76 |
| N3 sleep stage (min) | **84.21±37.23** | **45.71±37.88\*** |
| N3 sleep stage (%) | **21.70±7.83** | **9.90±7.97\*\*** |
| Non-rapid-eye movement (NREM) (min) | 321.00±78.85 | 383.86±111.29 |
| NREM (%) | 81.80±10.11 | 81.96±10.05 |
| Rapid-eye movement (REM) (min) | 82.04±54.54 | 76.83±29.52 |
| REM (%) | 18.20±10.11 | 18.04±10.05 |
| Sleep efficiency (%) | 71.13±18.87 | 81.52±16.78 |
| Sleep latency | **28.36±23.75** | **7.93±5.48\*\*** |
| Wake after sleep onset | 138.54±122.07 | 77.79±46.87 |
| Arousal time (min) | **165.71±118.75** | **84.31±48.77\*** |
| Arousal times | 16.21±8.06 | 12.50±10.70 |
| Arousal index (events/h) | 2.66±1.52 | 1.89±1.83 |
| Arousal time in REM | 39.43±28.30 | 66.67±39.80 |
| Arousal index in REM | **29.50±12.13** | **50.00±13.49\*\*\*** |
| Arousal time in NREM | **163.86±66.82** | **433.25±262.42\*\*** |
| Arousal index in NREM | **30.54±9.82** | **63.53±28.16\*\*** |
| Total sleep arousal times | **203.29±85.31** | **499.92±285.72\*\*** |
| Total sleep arousal index | **30.04±8.81** | **61.68±25.41\*\*** |
| Apnea-hypopnea index (events/h) | **5.88±3.43** | **52.53±29.78\*\*\*** |
| Apnea-hypopnea times | **42.00±25.31** | **430.58±294.38\*\*** |
| Obstructive apnea index (events/h) | **1.41±1.65** | **21.38±18.74\*\*** |
| Obstructive apnea times | **9.86±10.79** | **173.25±159.52\*\*** |
| Central apnea index (events/h) | **0.30±0.77** | **3.83±5.40\*** |
| Central apnea times | 2.07±5.14 | 35.50±54.65 |
| Mixed apnea index (events/h) | **0.05±0.09** | **6.19±9.08\*** |
| Mixed apnea times | 0.36±0.63 | 55.33±88.13 |
| Hypopnea index (events/h) | **4.11±2.65** | **21.17±16.88\*\*** |
| Hypopnea times | **29.71±21.00** | **166.50±155.08\*\*** |
| Longest apnea time (s) | **24.86±24.59** | **69.08±38.55\*\*** |
| Mean apnea-hypopnea duration (s) | **14.06±7.29** | **26.97±7.53\*\*\*** |
| Longest hypopnea time (s) | **60.29±28.51** | **99.75±24.15\*\*** |
| Average hypopnea time (s) | **27.91±6.86** | **35.74±8.24\*** |
| Oxygen desaturation index (events/h) | **5.48±3.66** | **49.73±29.24\*\*\*** |
| Lowest oxygen saturation (%) | **88.57±3.34** | **74.83±12.04\*\*\*** |
| Average oxygen saturation (%) | **95.00±1.79** | **92.50±2.81\*\*** |
| Longest oxygen desaturation (s) | 84.23±33.17 | 92.93±26.57 |
| Mean heart rate | **62.00±9.08** | **67.83±10.83\*** |
| Arrhythmia index (events/h) | 2.02±2.09 | 3.82±5.03 |
| Maximum heart rate | 94.00±4.60 | 103.08±18.70 |
| Minimum heart rate | 51.67±6.12 | 53.50±6.56 |
| Blood pressure elevation index (events/h) | 11.10±9.49 | 26.09±27.79 |
| The highest systolic blood pressure (mmHg) | 105.67±53.60 | 163.25±62.49 |
| Average systolic blood pressure (mmHg) | 80.83±40.36 | 117.58±41.10 |
| Average diastolic blood pressure (mmHg) | 62.83±31.35 | 84.25±29.45 |

### Cytokine analysis

There were not significantly different in IL-6 and TNF-α among three enterotypes patients (Figure 2).

**Figure 2.**
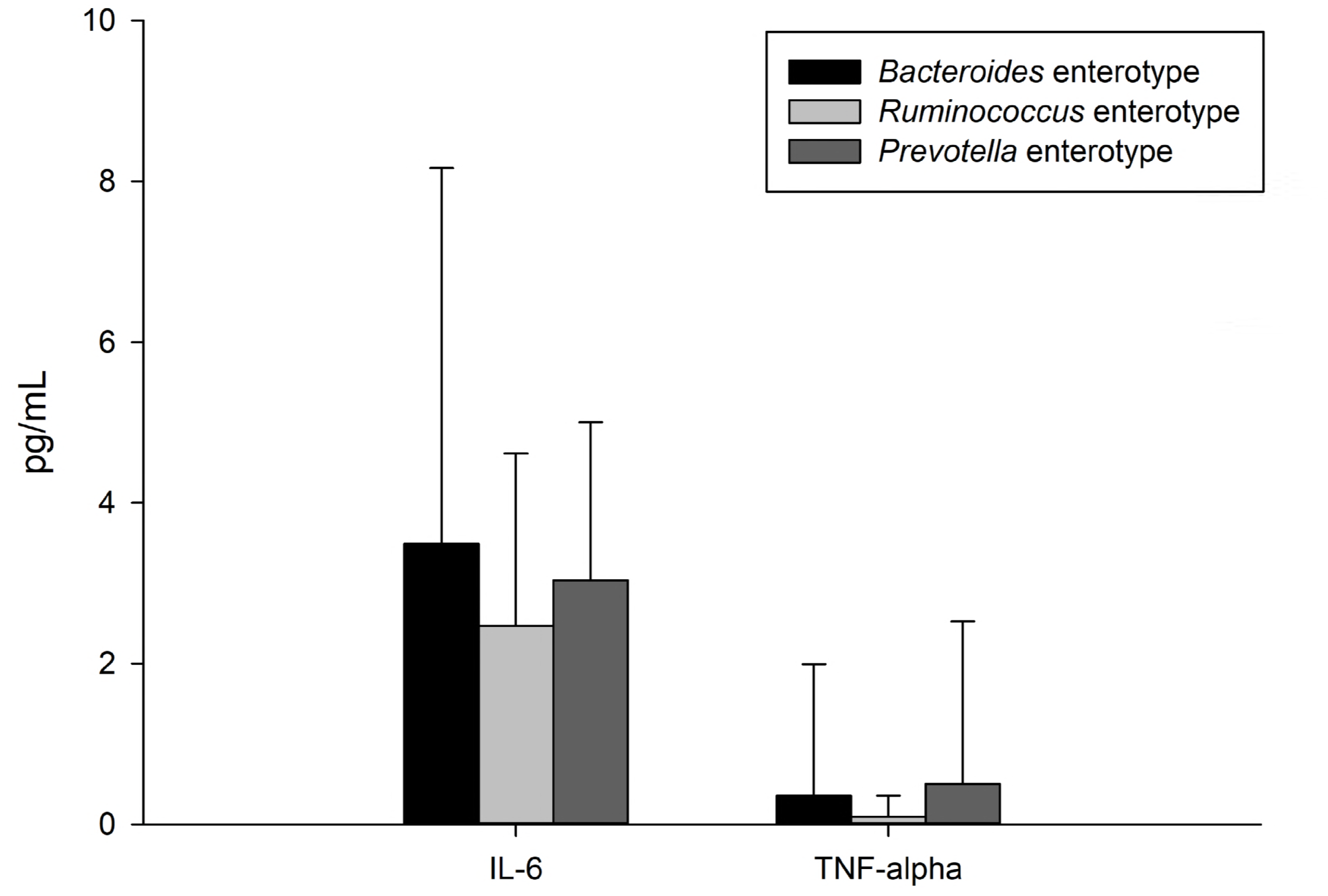
Cytokines levels analysis in three enterotypes subjects. IL: interleukin, TNF: tumor necrosis factor.

## Discussion

OSAHS is a systemic and comprehensive disorder associated with comorbidities, including cardiovascular disease, metabolic abnormalities, and neuropsychiatric and neurodegenerative disorders [4,5,7,9]. Thus, the IH mechanism alone is insufficient to interpret the complete pathogenesis of OSAHS because OSAHS is also affected by several other aspects, including the CNS. This study shows that central apnea indices are significantly elevated in AHI≥15 patients with the *Prevotella* enterotype, accompanied other parameters of disturbances in sleep. However, for AHI<15 results, changes in obstructive apnea indices and systolic BP are the remarkable observations in *Ruminococcus* enterotype and *Prevotella* enterotype subjects, respectively.

*Bacteroides* enterotype is associated with Western-style diets, including consuming high amounts of protein and fat. *Prevotella* enterotype is associated with diets high in carbohydrates (fiber) and simple sugars, whereas *Ruminococcus* species enterotype is linked to non-digestible carbohydrates [14]. Despite the fact that the *Bacteroides* predominant enterotype seems to be more common in IBD patients, the *Prevotella* enterotype is more representative in healthy subjects [15,16]. Our findings show that *Bacteroides* enterotype patients are not susceptible to OSAHS, in contrast to the susceptibility of *Prevotella* and *Ruminococcus* enterotype patients.

IH-exposed mice mimic OSAHS, causing profound alterations in gut microbiota. Hypoxia/re-oxygenation is the most pronounced [1], inducing an alteration in intestinal epithelial barrier markers and increasing intestinal permeability, leading to local and systemic inflammatory responses and consequent multi-organ morbidities [20,21]. However, only *Bacteroides* and *Prevotella* enterotypes can be classified in the rodent model, IH-exposed mice classify as the *Prevotella* enterotype [1], who is similar to our particularly OSAHS patients. It has also been shown that IH leads to gut microbiota alteration and accompanying endotoxin production [22]. It is that IH model creates an anoxic environment in the intestine, which is beneficial for obligate anaerobic bacterial growth, endogenous LPS production from gram-negative bacteria, and triggering inflammation. Notably, *Prevotella* is a genus of gram-negative anaerobic bacteria, and it tends to alter intestinal permeability [1,2]. Although this evidence only reveals the IH contribution to the pathogenesis, we speculated that SF is another principal contributor [3]. SF-induced mice manifest inflammation and enhanced production of endotoxins produced by gut microbiota, too [3]. In middle-aged nonobese males with OSAHS, disruption of the intestinal barrier and concurrent increased serum d-lactate levels possibly contribute to intestinal hyperpermeability and are significantly positively associated with IL-1β, IL-6, and TNF-α in serum [19] in which TNF-α elevation in *Prevotella* enterotype subjects is similar with our results, but it did not reach statistically significant differences. Moreover, LPS may play a key role in driving systemic inflammation, it has been shown in IH and SF modeling OSAHS models [1-3].

*Prevotella* enterotype patients with AHI≥15 in our results, suggesting that LPS production triggers downstream signaling pathways, leading to the subsequent release of pro-inflammatory IL-1β, IL-6, and TNF-α cytokines [23]. Furthermore, the elevation of LPS-binding protein [19] is also verified in OSAHS, mimicking rodent models [3] and patients [19], particularly regarding in the higher d-lactate level of OSAHS patients. Inflammatory mediators can be produced by peripheral and central cells. Peripheral inflammatory mediators may invade the CNS by crossing the blood– brain barrier, affecting behaviors and causing metabolic problems and psychiatric disorders [24]. Here, our data suggest that the gut microbiota impact the brain in OSAHS patients by modulating inflammatory responses. Additionally, we should mention that the *Prevotella* enterotype is linked to diets rich in simple sugars. Simple carbohydrate consumption has been hypothesized to be related to elevated BP values and obesity [25], as shown in our data. *Prevotella* enterotype patients had a higher BMI and hip circumference than *Bacteriodes* enterotype patients. Monosaccharide intake induces inflammation in epithelial cells and contributes to hypertension [25], linking to LPS production, which can stimulate systemic inflammatory cascades [1]. Inflammation mediates the pathogenesis of many physiological dysfunctions, such as metabolic syndrome and mental dysfunction [24], and thus might ultimately result in OSAHS-related metabolic comorbidities. Although *Ruminococcus* is associated with resistant starch, host health benefits from short chain fatty acids that have been demonstrated to regulate immune inflammatory responses [26]. The enriched bacteria *Ruminococcus* spp. and *Sutterella* spp. are found in autism spectrum disorder patients [27]. The abovementioned literature supports the hypothesis that microbiota disruption influences the pathophysiological process of OSAHS through a microbiota–gut–brain axis.

Although OSAHS is one of the most common sleep apnea syndromes (SAS), other types are mixed sleep apnea (MSA) and central sleep apnea (CSA). The prevalences of OSAHS, complex SAS (CompSAS), and central SAS are 84.0%, 15.0%, and 0.4%, respectively [28]. MSA generally describes the mixture of both obstructive and central apnea events during diagnostic sleep, although many central apnea index occurrence is also identified as MSA, which is sometimes referred to as CompSAS [29]. Whereas CompSAS is a form of CAS wherein the persistence or emergence of central apneas or hypopneas have disappeared with CPAP, patients have predominately obstructive or mixed apneas occurring at ≥5 events/h [30]. Additionally, reportedly, there is a high prevalence of hypertension and heart disease in patients with CompSAS [31]. In our data, the central apnea index and mixed apnea index were significantly increased in *Prevotella* enterotype patients with AHI≥15. Thus, abnormalities in electrocardiography, electroencephalography, electromyography, and electro-oculography results should be of more concern.

Both IH and SF have been shown to independently affect similar CNS regions in animal research [7]. The N1 sleep stage is associated with the transition from wakefulness to other sleep stages or the following arousal during sleep. A higher N1 percentage might mean more events of wakefulness and/or arousal, SF (episodic arousal from sleep), and sympathetic overactivity during sleep [10]. REM sleep dysregulation significantly contributes to cognitive distortions and dysfunctions that rely on emotion and memory functions are also affected [8]. Moreover, the effects of sleep deprivation on cognition have been investigated [32]. Thus, OSAHS patients have been found to have neurocognitive and emotional disorders, suggesting the modulation of various neurotransmitters during the sleep period [7]. Recently, a multicenter randomized controlled trial has been initiated evaluating the extent to which CPAP treatment improves neurocognitive dysfunction in OSAHS patients and examining the role of gut microbiota in this change [11]. Preliminary results suggest the viability of the hypothesis that microbiota modulate central nervous functions in OSAHS patients.

Although the neural mechanisms underlying SAS-induced brain injury have not been completely elucidated, repeated arousals enable the characterization of the different stages of sleep. In the present study, the N1 sleep stage, MAD, and arousal index were increased in *Prevotella* enterotype patients. BP was not significantly different among the three enterotype AHI≥15 patient groups, but mean diastolic pressure during sleep was >80 mmHg, which was similar to that observed in a previous study [10]. MDA can act as an indicator of the levels of sleep parameters and blood oxygenation for the evaluation of severe OSAHS patients [33]. When MAD is elevated, sleep apnea appears to be more likely to cause respiratory arousal and might impair sleep stability, resulting in SF. This outcome might then be that the transition of the N2 sleep stage (the longest stage of sleep) to the N3 sleep stage is a vulnerable period, which is interrupted in OSAHS patients, and the overall sleep pattern becomes light sleep [33]. Additionally, chronic SF induction elevates fat mass, alters fecal microbiota, promotes increased gut permeability, leads to systemic and adipose tissue inflammatory changes, and accompanies metabolic dysfunction [3]. These symptoms are known to be associated with OSAHS-related metabolic comorbidities, implying that the microbiota–gut–brain axis has a biaxial effect on the development of OSAHS pathology.

Contrastingly, evidence has shown that N1, N3, and REM sleep stages decrease and the N2 sleep stage increases in OSAHS patients [7]. However, a higher N1 percentage, a longer MAD, and a shortened REM sleep stage were revealed in AHI≥15 patients with OSAHS-induced hypertension [10,33]. Our findings reveal that BP plays a vital role, particularly for SAS, where BP is comprehensively regulated by the peripheral and central systems. Hence, future studies should re-examine these questions in subgroups of hypertensive and normotensive OSAHS patients to assess their general applicability.

The current study initiates a new approach to the study of sleep apnea through a combination of polysomnographic measurements with analysis of gut microbiota. Central apnea index, mixed apnea index, N1 sleep stage, MAD, and arousal indices were all increased in AHI≥15 patients with the *Prevotella* enterotype. Our results raise the possibility that the microbiota–gut–brain axis operates bidirectionally, with significant impact on the pathogenesis of OSAHS including functions of the gut and brain that eventually contribute to multiple end-organ morbidities.

## Materials and Methods

### Subjects

In total, 113 subjects were recruited and examined during a full night of PSG (SOMNOscreen™ plus PSG^+^; SOMNOmedics GmbH, Randersacker, Germany) by technologists in a sleep laboratory from 10 PM to 8 AM at the Department of Pulmonary and Critical Care Medicine. Fecal samples were collected the following morning. The Institutional Review Board of the Second Affiliated Hospital of Fujian Medical University approved this study (IRB No. 2017-78).

### OSAHS evaluation

All the subjects underwent PSG with a computerized polysomnographic system, simultaneously including electrocardiography, electroencephalography, electromyography, and electrooculography. After one night of examination, AHI were calculated as the total number of episodes of apnea (continuous cessation of airflow for at least 10 s) and hypopnea (reduction in airflow for ≥10 s with oxygen desaturation ≥4%) by dividing the total sleep by events, according to the diagnostic criteria of the American Academy of Sleep Medicine. AHI<15 events/h was defined as non-OSAHS and AHI≥15 events/h as OSAHS in this study, as reported previously [10,19].

### Cytokine analysis

IL-6 and TNF-α were assayed by BD Human Enhanced Sensitivity Cytometric Bead Array Kit (BD Biosciences, New Jersey, USA) as described previously [18]. The standard coefficient of determination (r^2^) was greater than 0.995.

### Sampling, DNA extraction, and 16S rRNA gene amplification sequencing

Samples were collected and stored in a Microbiome Test Kit (G-BIO Biotech, Inc., Hangzhou, China). Magnetic bead isolation was performed to extract genomic DNA using a TIANamp stool DNA kit (TIANGEN Biotech Co., Ltd., Beijing, China), according to the manufacturer’s instructions. The concentration of extracted DNA was determined by a Nanodrop ND-1000 spectrophotometer (Thermo Electron Corporation, USA), and DNA quality was confirmed using 1.0% agarose gel electrophoresis with 0.5 mg/mL ethidium bromide.

Isolated fecal DNA was used as a template to amplify the V3 and V4 hypervariable regions of the bacterial 16S ribosomal RNA gene. The V3 and V4 regions were PCR-amplified (forward primer, 5′-ACTCCTACGGGAGGCAGCAG-3′; reverse primer, 5′-GGACTACHVGGGTWTCTAAT-3′). The 16S target-specific sequence contained adaptor sequences permitting uniform amplification of a highly complex library ready for downstream next-generation sequencing on Illumina MiSeq (Illumina, USA). Negative DNA extraction controls (lysis buffer and kit reagents only) were amplified and sequenced as contamination controls. The amplicons were normalized, pooled, and sequenced on the Illumina MiSeq platform using a V3 reagent kit with 2 × 300 cycles per sample and with imported and prepared routine data (samsheet) run in the MiSeq sequence program. After sequencing, Q30 scores were ≥70%, the percentage of clusters passing filter (i.e., cluster PF) was ≥80%, and there were at least 30,000 clean tags. Finally, image analysis and base calling were conducted with MiSeq Control Software.

### Bioinformatic, predictive function and statistical analyses

Based on the Quantitative Insights into Microbial Ecology bio-informatic pipeline for performing taxonomy assignment by the operational taxonomic unit method, we used data of 113 sequences to analyze the fecal microbiota taxa. We analyzed differences in gut microbiota using the Wilcoxon test, as appropriate, and performed principal coordinate analysis on the basis of the Bray–Curtis distance function, using R statistics. We performed other analyses using statistically with SPSS version 19.0 (SPSS Inc., Chicago, IL, USA); data were analyzed by *t*-test or one-way ANOVA, followed by Scheffe post hoc analyses. We considered a two-sided *p* value of <0.05 to be statistically significant.

## Declarations

### Conflict of interest

The authors declare that they have no financial and personal relationships with others that may inappropriately influence the results and interpretation in this manuscript.

### Role of the funding source

None.

### Contributors

Conception and design: CYK, HPZ, YMZ

Acquisition of data: CYK, AKH, JMF, LMH, JHY, HZS

Analysis and interpretation of data: CYK, AKH, JMF, HPZ, YMZ Drafting or revising of the article: CYK, HPZ, YMZ

Final approval of the manuscript: All authors read and approved the final manuscript

## Acknowledgments

We thank all the participants and their family who took part in this study. The authors appreciate for Huan Wu (G-BIO Biotech, Inc., Hangzhou, China) assist us to carry out the bioinformatic analysis. The authors would like to thank the Fujian Provincial Health and Family Planning Commission, China, under contract No. 2018-CX-36; the Fujian Province Science and Technology Project, China, under contract No. 2018J01290; the Quanzhou Science and Technology Project, China, under contract No. 2017Z016 and No. 2018Z107. This article was also subsidized by academic funding from the Second Affiliated Hospital of Fujian Medical University (serial No. BSH001).

